# Impact of library preparation-induced low GC sequencing bias on *Salmonella* serotype prediction using SeqSero2

**DOI:** 10.1101/2020.03.20.997510

**Authors:** Shaoting Li, Shaokang Zhang, Xiangyu Deng

## Abstract

A recent study by Uezle et al. reported a lack of O antigen prediction by SeqSero2. We found that Nextera XT-prepared genomes in the study were abnormally challenging for O antigen prediction because of unusually high sequencing bias against low GC regions that was not seen in similarly prepared genomes from other laboratories. We recommend that SeqSero2 users be mindful of the GC-related sequencing bias when analyzing Nextera XT-prepared genomes. Although it is unusual for such biases to compromise serotype prediction by SeqSero2 per our knowledge and analysis, it is unknown whether they could affect subtyping and characterization of other low-GC regions, such as *Salmonella* pathogenicity islands, when genome assembly is affected by such biases.

---

SeqSero2 (1) and its predecessor SeqSero (2) predict *Salmonella* serotypes from whole genome sequencing (WGS) data by targeting genetic determinants of serotype without resorting to surrogate markers, such as MLST. This approach maintains continuity with the well-established scheme for phenotypic serotypes but may generate incomplete prediction of an antigenic profile should a serotype determinant gene be poorly sequenced by WGS (2).

DNA libraries prepared by the Illumina Nextera XT kits are known to produce suboptimal sequencing coverage at low-GC regions; this bias has implications for subtyping and metagenomics analyses (3-6). The lipopolysaccharide O antigen determinants of *Salmonella* in the *rfb* gene cluster feature considerably lower GC content (∼30%) than the genome-wide GC average of *Salmonella* (∼52%). A recent evaluation of *Salmonella* serotype prediction tools by Uezle et al. reported a lack of O antigen prediction by SeqSero2 (7). The authors convincingly attributed such predictions to library preparation-induced low GC sequencing bias caused by the Nextera XT kits. In comparison, genomes prepared by the newer Illumina Nextera Flex kits were free of the issue (7).

Lack of O antigen prediction in the Uezle study was alarmingly prevalent, which prompted us to reanalyze their data to investigate the cause of the reported issue. Compared to a representative set of *Salmonella* genomes from public health laboratories in United States (U.S.) and England, Nextera XT-prepared genomes in the Uezle study appeared to be disproportionately overrepresented by predictions that lacked an O antigen call (Table 1), the vast of which belonged to the serogroup O7 (Table 2). These O7 genomes as well as Nextera XT-prepared genomes of other common serogroups (O4, O8, and O9) in the Uezle study were significantly more susceptible to GC content-associated sequencing bias against low GC regions (Figures 1 and 2). The biases were significant enough to affect *de novo* genome assembly as measured by the L50 score (Figure 2) and likely contributed to the uncharacteristically low antigen prediction accuracy by SISTR, another tool evaluated in the Uezle study (34.5% full match rate vs. 41.9% in a previous evaluation (8)). In our benchmark dataset, genomes from U.S. NARMS were prepared by Illumina TruSeq kits and least affected by the sequencing bias (Figure 1). These genomes were used in the previous evaluation of SeqSero2 (1); their bias-free nature (Figure 3) may explain the discrepant results between the previous and the Uezle studies, particularly the performance of the micro-assembly workflow that requires sufficient sequencing coverage of the *rfb* region to assemble O antigen determinant genes.

**Table 1.** Prevalence of serotype predictions that lack an O antigen call by the microassembly workflow of SeqSero2 using *Salmonella* genomes from different public health laboratories

| Dataset | Total | library preparation | Predictions that lack an O antigen call | % Predictions that lack an O antigen call |
| --- | --- | --- | --- | --- |
| BfR<sup>a</sup> | 578 | Nextera XT | 71 | 12.3% |
| US FDA<sup>b</sup> | 3929 | Nextera XT | 33 | 0.8% |
| US PulseNet<sup>c</sup> | 196 | Nextera XT | 5 | 2.6% |
| PHE<sup>d</sup> | 202 | Nextera XT | 0 | 0 |
| US NARMS<sup>e</sup> | 2280 | TrueSeq | 5 | 0.2% |

<sup>a</sup>Genomes ( $n=1263$ ) from animal production, food and environment in Germany under the BioProject PRJEB31846 were analyzed. Out of the 1263 genomes, 578 were prepared by Nextera XT kits, of which 71 were missing an O antigen call. Another 685 were prepared by Nextera Flex kits, of which 3 were missing an O antigen call. NCBI accession numbers can be found at http://www.denglab.info/SeqSero2/AEM_letter_datasets.xlsx.
<sup>b</sup>Genomes ( $n=3929$ ) used by FDA for an evaluation study of SeqSero2 (unpublished). NCBI accession numbers can be found at http://www.denglab.info/SeqSero2/AEM_letter_datasets.xlsx.
<sup>c</sup>Genomes ( $n=196$ ) sequenced by state and local health departments in the US for national surveillance of *Salmonella*. Genomes were randomly selected from BioProject PRJNA230403 to represent 16 major serotypes, including serotype Braenderup, Infantis, Montevideo, Thompson, Agona, Heidelberg, Saintpaul, Typhimurium, Hadar, Kentucky, Muenchen, Newport, Berta, Enteritidis, Javiana, and Panama. NCBI accession numbers can be found at http://www.denglab.info/SeqSero2/AEM_letter_datasets.xlsx.
<sup>d</sup>Genomes ( $n=202$ ) were randomly selected from the Public Health England BioProject PRJNA248792 to represent 16 major serotypes as aforementioned. NCBI accession numbers can be found at http://www.denglab.info/SeqSero2/AEM_letter_datasets.xlsx. Genomes were prepared by Nextera XT kits according to the annotation of WGS data in the depository.
<sup>e</sup>Genomes ( $n=2280$ ) from human clinical isolates submitted to CDC NARMS in 2015 (1). NARMS performs surveillance for antimicrobial resistance in *Salmonella* (https://www.cdc.gov/narms/index.html); every 20th isolate, along with serotype information, is submitted by state and local health departments in the US. NCBI accession numbers can be found at http://www.denglab.info/SeqSero2/AEM_letter_datasets.xlsx.

**Table 2.** Summary of serotype predictions that lack an O antigen call by the microassembly workflow of SeqSero2 among Nextera XT-prepared genomes in the Uezle study

| Serotypes<sup>a</sup> | Total | Predictions that lack an O antigen call | O group | % O antigen less predictions |
| --- | --- | --- | --- | --- |
| Virchow | 3 | 3 | O7 | 100.0% |
| Bareilly | 4 | 3 | O7 | 75.0% |
| Infantis | 69 | 29 | O7 | 42.0% |
| Mbandaka | 58 | 17 | O7 | 29.3% |
| Paratyphi B var. Java | 55 | 5 | O4 | 9.1% |
| Agona | 41 | 3 | O4 | 7.3% |
| Typhimurium | 61 | 4 | O4 | 6.6% |

<sup>a</sup>Only serotypes with at least 3 genomes that produced predictions without an O antigen call are shown. These genomes accounted for 90.1% of predictions that lacked an O antigen call from *Salmonella* isolates from animal production, food and environment in Germany ( $n=1263$ ) under the BioProject PRJEB31846.

**Figure 1.**
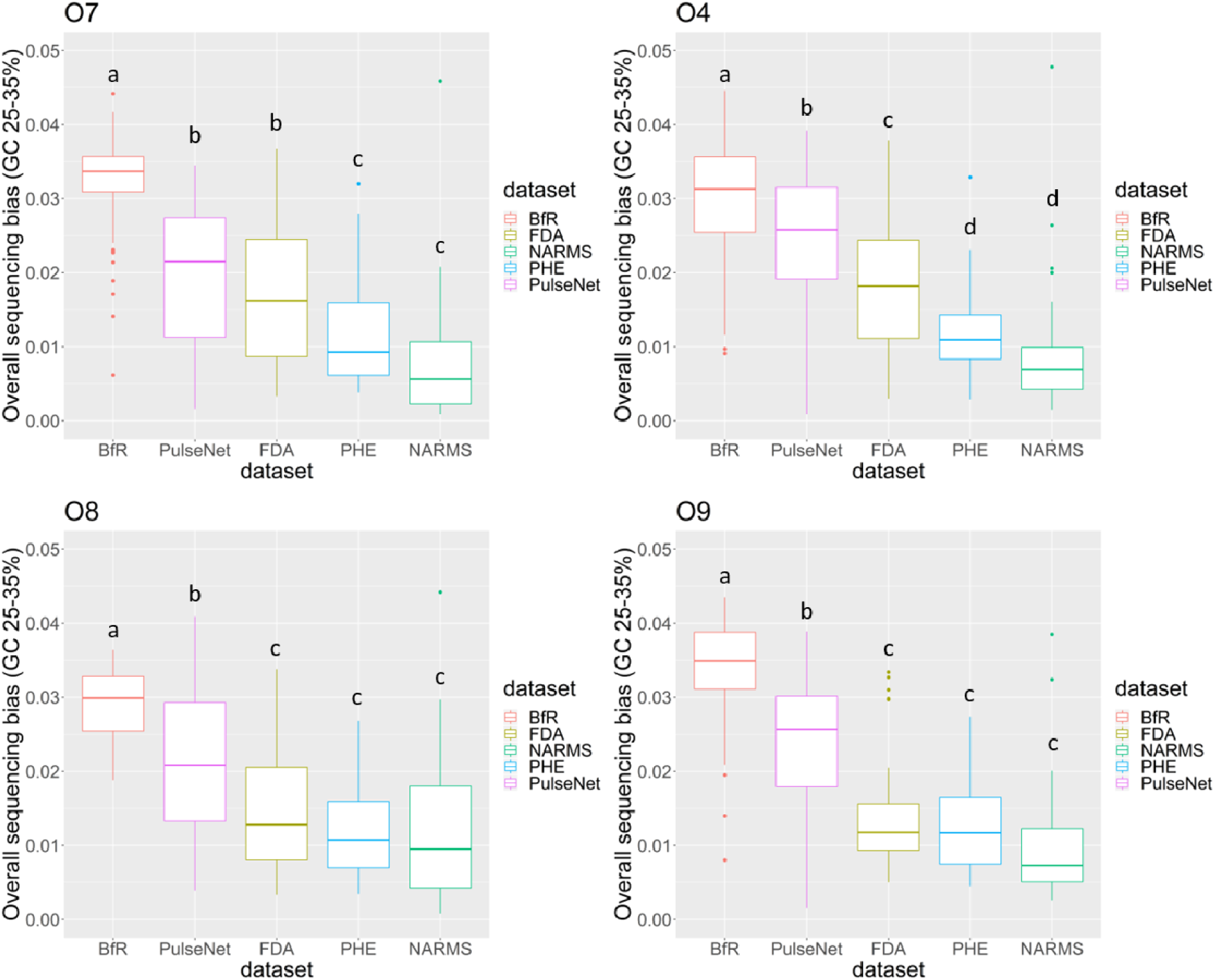
Comparison of GC-associated sequencing biases of low GC regions in genomes from four major O antigen groups. Low GC regions are defined as regions of GC content between 25% and 35%, which is similar to the GC content of O antigen determinant genes (*wzx/wzy*). GC-associated sequencing bias was calculated according to methods described in (3). *Salmonella* genomes from five datasets were analyzed, including BfR (*n*=415), PulseNet (*n*=158), FDA (*n*=145), PHE (*n*=160) and NARMS (*n*=155). Genomes of four major O antigen groups (O7, O4, O8 and O9) in these datasets were analyzed. For the PulseNet, FDA, PHE and NARMS datasets, the O7 group includes serotype Braenderup, Infantis, Montevideo and Thompson; the O4 group includes serotype Agona, Heidelberg, Saintpaul and Typhimurium; the O8 group includes serotype Hadar, Kentucky, Muenchen and Newport; and the O9 group includes serotype Berta, Enteritidis, Javiana and Panama. Up to 10 genomes of each serotype were randomly selected from each dataset. For Nextera XT-prepared genomes in the Uezle study (*n*=578), all the O7, O4, O8, and O9 genomes were included except serotype Enteritidis. This serotype was overrepresented in the Uezle study (*n*=115) and 20 genomes were randomly selected for this analysis. Different characters on the top of boxes indicate there is significant difference (*P* < 0.05, ANOVA) in between.

**Figure 2.**
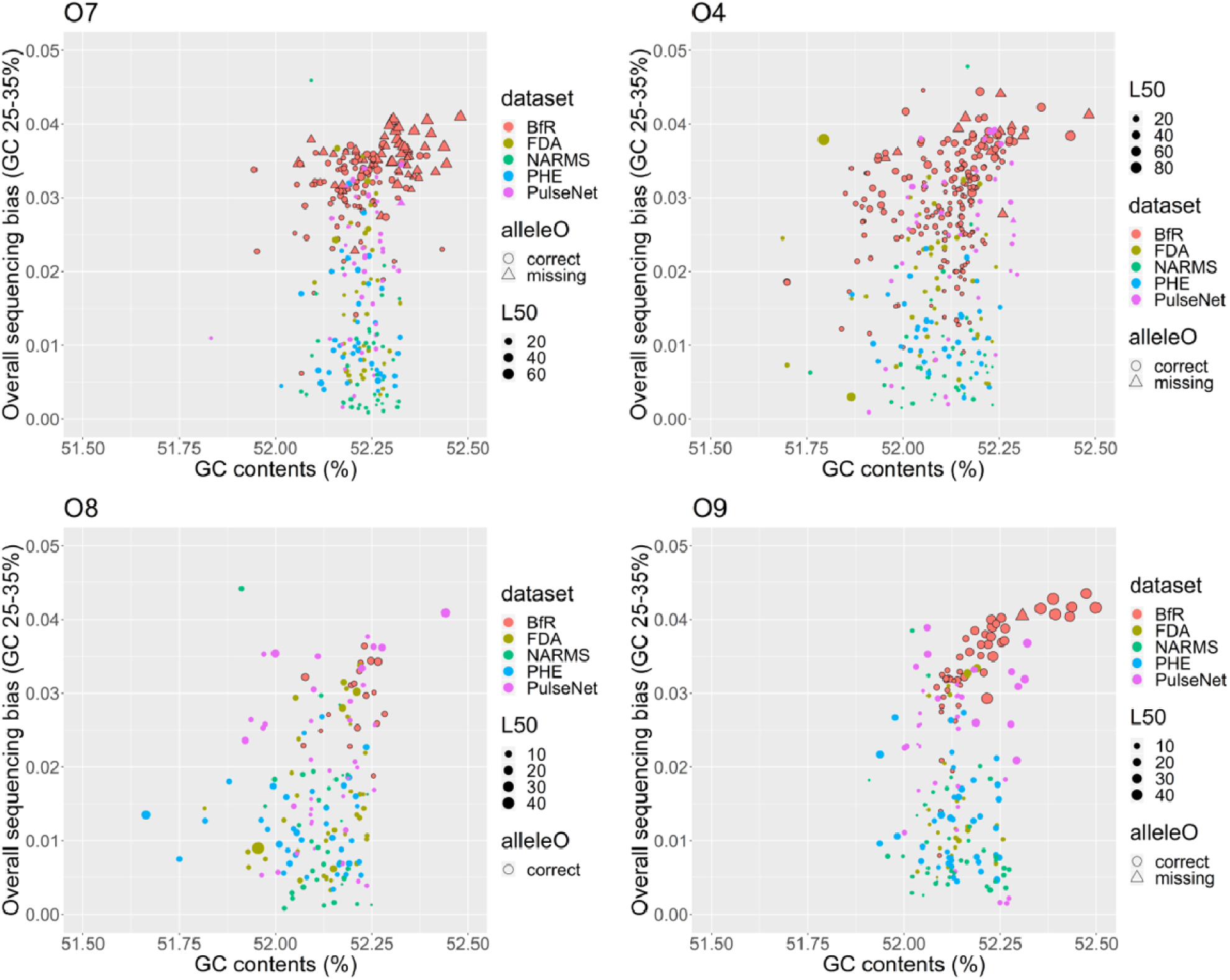
Sequencing biases of low GC regions and assembly quality of entire genomes. Low GC regions are defined as regions of GC content between 25% and 35%, which is similar to the GC content of O antigen determinant genes (*wzx/wzy*). Genome assembly quality is represented by L50. GC-associated sequencing bias was calculated according to (3). *Salmonella* genomes from five datasets were analyzed, including BfR (*n*=415), PulseNet (*n*=158), FDA (*n*=145), PHE (*n*=160) and NARMS (*n*=155). Serotypes of four major O antigen groups (O7, O4, O8 and O9) in these datasets were analyzed separately. The O7 group includes serotype Braenderup, Infantis, Montevideo and Thompson; the O4 group includes serotype Agona, Heidelberg, Saintpaul and Typhimurium; the O8 group includes serotype Hadar, Kentucky, Muenchen and Newport; and the O9 group includes serotype Berta, Enteritidis, Javiana and Panama. Up to 10 genomes of each serotype were randomly selected from each dataset. For Nextera XT-prepared genomes in the Uezle study (*n*=578), all the O7, O4, O8, and O9 genomes were included except serotype Enteritidis (*n*=115). This serotype was overrepresented in the Uezle study and 20 genomes were randomly selected for this analysis.

**Figure 3.**
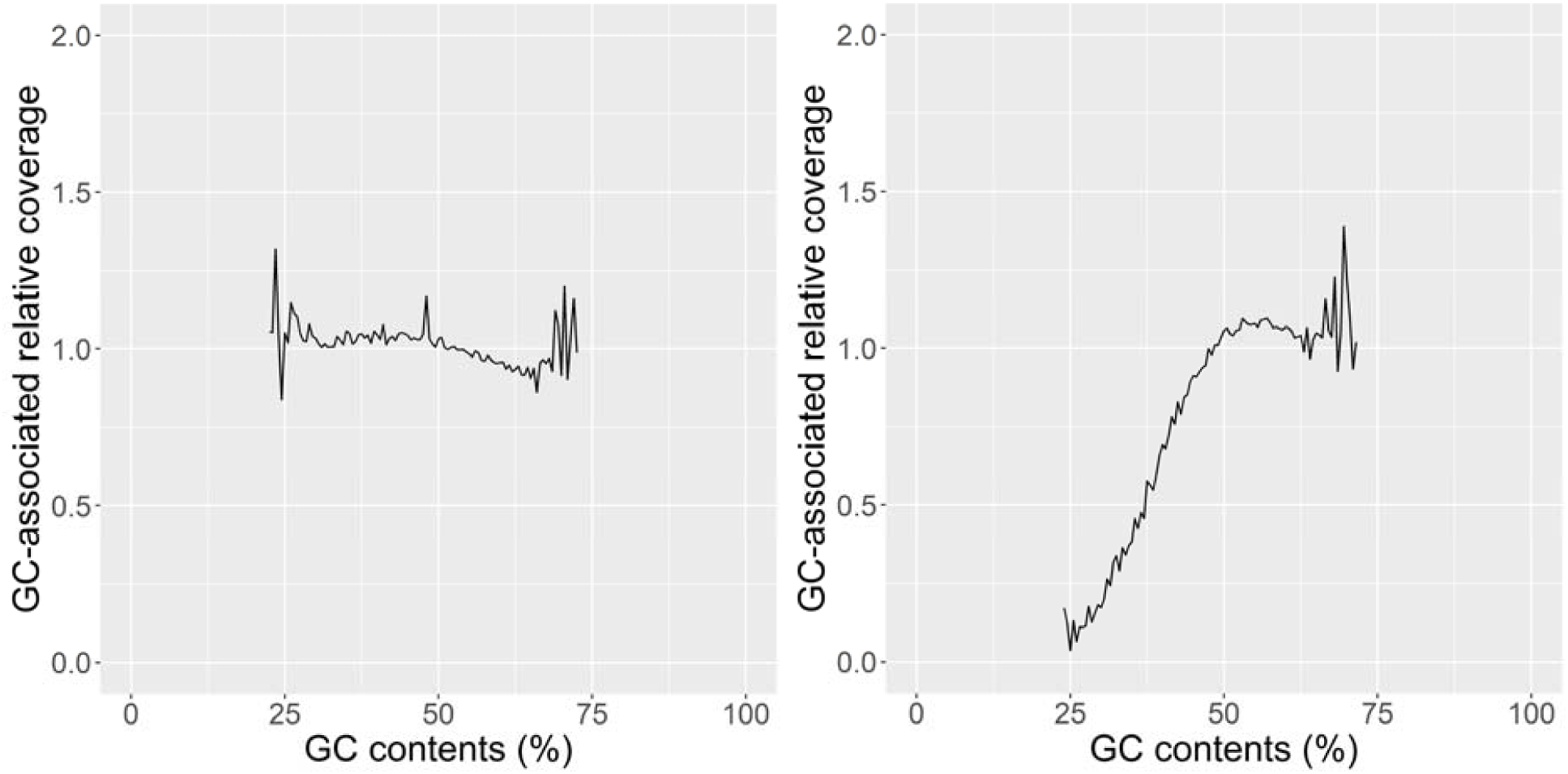
An example of sequencing bias profile by TruSeq (left, accession number SRR5740049) and Nextera XT (right, accession number ERR3581017) prepared genomes. The analyzed isolates were all serotype Infantis genomes. GC-associated relative coverage was calculated according to methods described in (3).

While unrelated to sequencing bias, the Uezle study reported misidentification of serotype Enteritidis as serotype Hillingdon, due to a misidentification of serogroup O9 as O9,46 that was specific to the *k*-mer workflow of SeqSero2. This issue was independently identified by multiple laboratories in the U.S. and addressed in later releases of SeqSero2. We noted that the Uezle study described SeqSero2 workflows with obsolete terms such as “*k*-mer mode” and “allele-mode” and did not mention which version of SeqSero2 was evaluated. These terms were used only in the earliest test release of SeqSero2 prior to the first stable version (v1.0.0) that was published (1).

In conclusion, the genomes used in the Uezle study were abnormally challenging for O antigen prediction because of unusually high sequencing bias that was not seen in similarly prepared genomes from other laboratories. We recommend that SeqSero2 users be mindful of the GC-related sequencing bias when analyzing Nextera XT-prepared genomes. Although it is unusual for such biases to compromise serotype prediction by SeqSero2 per our knowledge and analysis, it is unknown whether they could affect subtyping and characterization of other low-GC regions, such as *Salmonella* pathogenicity islands (9, 10) when genome assembly is affected by such biases (Figure 2).

## Acknowledgments

We thank Patricia Fields, Blake Dinsmore, Ana Lauer and Jessica Chen of US CDC and Ruth Timme, Shaohua Zhao and Sunee Himathongkham of US FDA for providing WGS data and/or helpful discussion.

## Notes

http://denglab.info/static/AEM_letter_datasets.xlsx

